# CypA is a molecular barrier to HIV emergence from Eastern chimpanzees

**DOI:** 10.64898/2026.09.08.747897

**Authors:** Michael J. Young, Fabricio Cotera, Clare C. C. Gill, Jacob A. Lewis, Frank G. Whitby, Alaa Abdellatif, Joana L. Rocha, Michael Singer, Mike A. Nalls, Faraz Faghri, Peter H. Sudmant, Owen Pornillos, Molly Ohainle

## Abstract

Of the two lineages of lentivirus that endemically infect chimpanzees, only one has served as a source of infections in humans. Using HIV-CRISPR screening, we identify factors that limit the replication of lentivirus from Eastern chimpanzees. We find that CypA restricts nonzoonotic Eastern chimpanzee lentivirus due to a unique feature encoded in capsid. The ability of CypA to restrict lentivirus replication is surprising given that CypA is a host factor exploited by HIV to promote its replication. The selective ability of CypA to restrict this virus provides a molecular explanation as to why lentivirus from Eastern Chimpanzees has never been observed to infect humans. Our work has important implications for understanding the origins of HIV-1 and the potential for novel zoonosis of primate lentiviruses.

## Introduction

A virus must overcome numerous ecological, immunological and molecular barriers to successfully cross to humans and become a human pathogen [1,2]. The massive, ongoing human immunodeficiency virus pandemic (HIV-1 M), originated with a cross-species transmission event from chimpanzees to humans a century ago [3–5]. Of the four subspecies of chimpanzee, each with distinct geographic ranges (Fig. 1A), two chimpanzee subspecies, Central (*Pan troglodytes troglodytes* or *Ptt*) and Eastern (*Pan troglodytes schweinfurthii or Pts)*, are endemically infected with distinct HIV-related simian immunodeficiency viruses (SIVs) [6]. SIV transmission to humans, while rare, has occurred on multiple occasions [7,8]. Curiously, only SIV from Central chimpanzees (SIVcpz*Ptt*) is the source of all HIV-1 infections in humans, including pandemic HIV-1 M (Fig. 1B) as well as other non-pandemic HIV-1 infections [3]. SIV from Eastern chimpanzees (SIVcpz*Pts*) has never been observed to infect humans despite similar SIV prevalence and extensive primate-human interaction [9]. We hypothesize that suboptimal molecular interactions between host and virus may serve as a barrier to SIVcpz zoonosis from Eastern but not Central chimpanzees.

**Figure 1.**
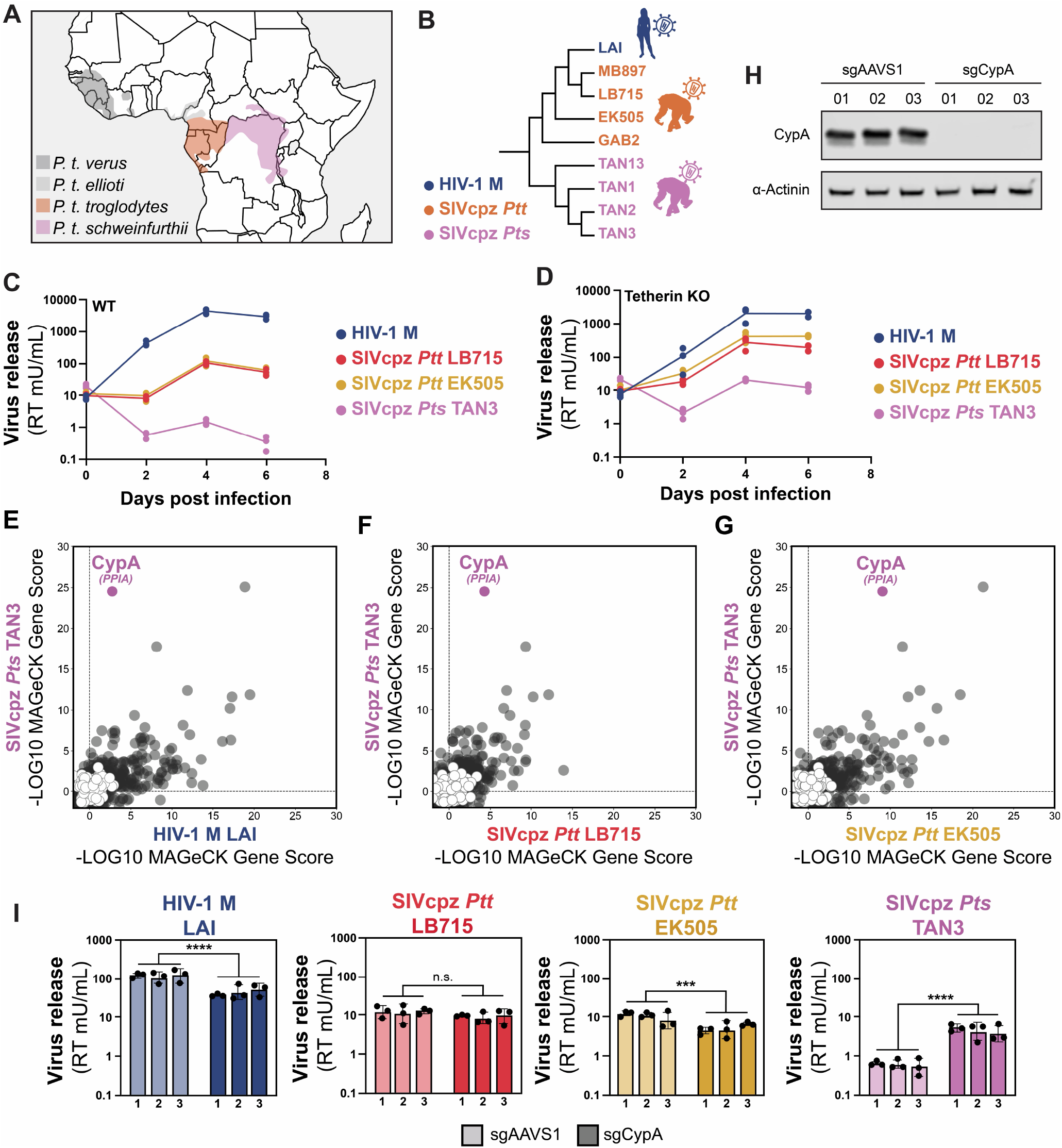
HIV-CRISPR screening identifies CypA as a restriction factor against SIVcpz*Pts*. **(A)** Geographic distribution of the 4 subspecies of chimpanzee including Eastern (*P. t. schweinfurthii*) and Central (*P. t. troglodytes*) chimpanzees. **(B)** PhyML Maximum-likelihood cladogram of full-length SIVcpz and HIV-1 M nucleotide genomes. SIVcpz*Pts* in Purple; SIVcpz*Ptt* in Orange; HIV-1 M in Blue. **(C)** Virus produced in THP-1 cells infected with input-normalized VSV-G pseudotyped HIV-1 and SIVcpz viruses across 2-6 days post infection (dpi). Virus was measured using the SG-PERT assay to quantify reverse transcriptase activity in viral supernatants (mU/mL RT activity). **(D)** Same as C but in THP-1 cells edited to lack human Tetherin. (**E** to **G**) Zoonosis Implicated sgRNA Assembly (ZIRA) THP-1 HIV-CRISPR screens. −log10 MAGeCK Gene Scores (positive scores only) for HIV-1 M LAI, SIVcpz*Ptt* LB715 and EK505 and SIVcpz*Pts* TAN3 in Tetherin KO THP-1 cells with the ZIRA library. SIVcpz*Pts* TAN3 (Y-axis in **E, F** and **G**) compared to HIV-1 M LAI (X-axis in **E**), SIVcpz*Ptt* LB715 (X-axis in **F**) and SIVcpz*Ptt* EK505 (X-axis in **G**). Non-Targeting Control (NTC) scores are shown in white; all other genes in grey. Source data for **E** to **G** can be found in Data S4 and S5. **(H)** CypA Western blot in negative controls (sgAAVS1) and CypA knockout THP-1 cell pools (sgCypA). Blots were probed for CypA and α-Actinin as a loading control. **(I)** THP-1 control (sg*AAVS1*) and CypA (sgCypA) knockout pools infected with input-normalized HIV-1 M LAI (blue), SIVcpz*Ptt* LB715 (red), SIVcpz*Ptt* EK505 (yellow) and SIVcpz*Pts* TAN3 (purple). Production of viral particles was measured using the SG-PERT secreted reverse transcriptase assay at three days post-infection. Data are presented as mean ± SD of triplicates from a single representative experiment. Statistical significance was determined by a two-tailed, unpaired Student’s t-test. *P<0.05; **P<0.01; ***P<0.001; ****P<0.0001; n.s.= not significant.

Successful zoonosis requires a virus to simultaneously evade antiviral recognition while maintaining optimal host factor engagement. Antiviral factors in primates are rapidly-evolving due to long-term selection driven by lentiviruses, retroviruses and related retroelements [10,11]. Antagonism of the genes Tetherin (*BST2*) and APOBEC3 were key steps in the adaptation of SIVcpz to successfully emerge as a pandemic in humans [12–14]. By contrast, the host protein Cyclophilin A (CypA or *PPIA*) and the cyclophilin domain-containing nuclear pore protein RANBP2 are important “dependency” factors required by HIV-1 for replication in host cells [15,16].

High throughput genetic screens can identify host factors that play important roles in HIV replication [17–19]. We developed a novel virus packageable screening approach, HIV-CRISPR, that is uniquely powerful in successfully identifying both known and novel lentiviral host factors across a range of viral and host cell systems [17,20,21]. In this study, we test the hypothesis that human cells encode host cell factors that engage differentially with distinct chimpanzee lentiviruses and present unique molecular barriers to zoonosis. We perform HIV-CRISPR screening and identify all genes important for the replication of SIVcpz viruses from both Eastern and Central chimpanzees in human cells. Whole-genome and sublibrary screens unexpectedly identify CypA as a key restrictor of Eastern chimpanzee SIVcpz (SIVcpz*Pts*) in human cells. We identify a unique feature in SIVcpz*Pts* capsid sequences that leads to a block early in the viral lifecycle before integration of viral genomes. CypA restriction is a conserved feature across viruses found in Eastern but not Central chimpanzees. Our results are the first to identify a block unique to SIV from Eastern chimpanzees. We propose that CypA serves as a key molecular barrier to SIVcpz zoonosis from Eastern chimpanzees to humans.

## Results

### CypA is a specific restrictor of nonzoonotic SIVcpz

To understand why SIVcpz from Eastern chimpanzees has never crossed into humans, we first compared replication of HIV-1 and SIVcpz viruses in human immune cells. Central chimpanzee lentiviruses (SIVcpz*Ptt* strains LB715 and EK505) replicate less well compared to the prototype HIV-1 M virus, LAI (Fig. 1C). The Eastern chimpanzee lentivirus (SIVcpz*Pts* TAN3) replicates even less efficiently (Fig. 1C). Knockout of human Tetherin (fig. S1), a known molecular barrier to SIVcpz replication in human cells [12], increases infection of all SIVcpz viruses but importantly replication of SIVcpz*Pts* TAN3 remains limited (Fig. 1D). We used genome-wide HIV-CRISPR screening [17,22] to define all cell-autonomous host factors that contribute to the relatively poor replication characteristic of the Eastern chimpanzee SIVcpz*Pts* lentivirus. HIV-CRISPR screening assays for effects of gene knockouts on HIV replication by quantifying HIV-CRISPR genomes packaged into released virions after infections of pools of HIV-CRISPR knock-out cells (fig. S2A) [17]. Our genome-wide HIV-CRISPR screens identify host factors involved in both HIV-1 and SIVcpz replication (fig. S2, B-D; Data S1 and S2). To enable open exploration of data in this study and comparison to previously published HIV-CRISPR screen data, we built an open data commons, CRISPRvirus (www.CRISPRvirus.org) (fig. S3). As expected, host factors known to be important for lentiviral transcription, including *CCNT1, NFKB1, EP300* and *MED26* [23–26], are depleted in all screens (fig. S2E). Conversely, non-targeting control (NTC) guides and guides targeting negative control antiviral genes are neither enriched nor depleted (fig. S2F). Type I Interferon pathway genes including *IFNAR2, STAT2* and *IRF9* are enriched in all screens (fig. S2, D and G), likely reflecting either tonic IFN signaling in THP-1 cells or sensing of viral infection that leads to ISG induction. Endosomal trafficking-related genes, including *C16orf62* and *COMMD3*, score strongly as restrictors of all viruses (fig. S2, D and H). However, these genes were not enriched in a screen with HIV-1 LAI encoding the native HIV-1 envelope, suggesting they are specific to the endosomal entry pathway of VSV-G (fig. S4). Tetherin (*BST2*), a known inhibitor of SIVcpz viruses in human cells, shows enrichment for all SIVcpz viruses but not HIV-1 M LAI (fig. S2, D and I). Genes involved in the biosynthesis of glycophosphatidylinositol (GPI) anchors (*GPAA1* and *PIGU*) [27] are also strongly enriched (fig. S2, D and I) consistent with the requirement of GPI anchoring in Tetherin antiviral activity [13]. To more robustly define the key genes important specifically for SIVcpz*Pts* TAN3 replication, we designed the <u>Z</u>oonosis-Implicated sg<u>R</u>NA <u>A</u>ssembly ZIRA) to target the strongest hits from our initial screen (fig. S5A; Data S3). We enriched specifically for Tetherin-independent restriction factors by performing ZIRA HIV-CRISPR screens in Tetherin knockout cells (fig. S5, B to E; Data S4 and S5). Remarkably, screening with the ZIRA library identifies a single gene, Cyclophilin A (*PPIA*), as the sole, specific restrictor of the SIVcpz*Pts* TAN3 virus as compared to all other viruses (Fig. 1, E, F and G). Infection of CypA knockout THP-1 cells (Fig. 1H) shows ~7-fold rescue for the SIVcpz*Pts* TAN3 virus compared to negative control cells (*AAVS1*) (Fig. 1I). Therefore, CypA is a specific restrictor of the nonzoonotic Eastern chimpanzee SIV.

### CypA restriction is dependent on capsid

CypA is a host factor that facilitates HIV-1 replication by stabilizing the capsid to allow for proper timing of uncoating and by protecting the capsid from host innate immune recognition [20,28–32]. Therefore, it is surprising that CypA restricts Eastern chimpanzee lentiviruses (SIVcpz*Pts*). To ask if the capsid (CA) is the molecular determinant of CypA restriction, we generated an isogenic set of chimeric GFP-reporter viruses harboring different SIV capsids in an otherwise HIV-1 backbone (Fig. 2A). Infection of CypA knockout THP-1 cells shows a ~4-fold increase in infection for only the SIVcpz*Pts* TAN3 capsid-encoding virus (Fig. 2B). To ask if CypA restricts other Eastern chimpanzee SIVcpz*Pts* capsid sequences, we tested additional SIVcpz*Pts* capsids in parallel with additional Central chimpanzee SIV capsids (Fig. 1B and Fig. 2C). CypA knockout leads to an increase in infection for all Eastern chimpanzee SIVcpz*Pts* capsids tested (TAN1, TAN2, TAN13) but does not rescue any of the Central chimpanzee SIVcpz*Ptt* capsid encoding viruses (MB897, GAB2) (Fig. 2C). Consistent with the CypA knockout data, infection in the presence of CsA, an inhibitor of CypA, increases infection with SIVcpz*Pts* capsid-encoding viruses (TAN3 and TAN13) but not SIVcpz*Ptt* or HIV-1 capsids (Fig. 2D). Blocking capsid-CypA interaction with CsA rescues infection of SIVcpz*Pts* capsids across a panel of immune cell lines (fig. S6A). The overall restriction phenotype measured across cell lines is not strictly correlated with levels of CypA (fig. S6, B and C), suggesting that CypA restriction is not determined strictly by steady-state expression levels and instead depends on other aspects of capsid-CypA biology.

**Figure 2.**
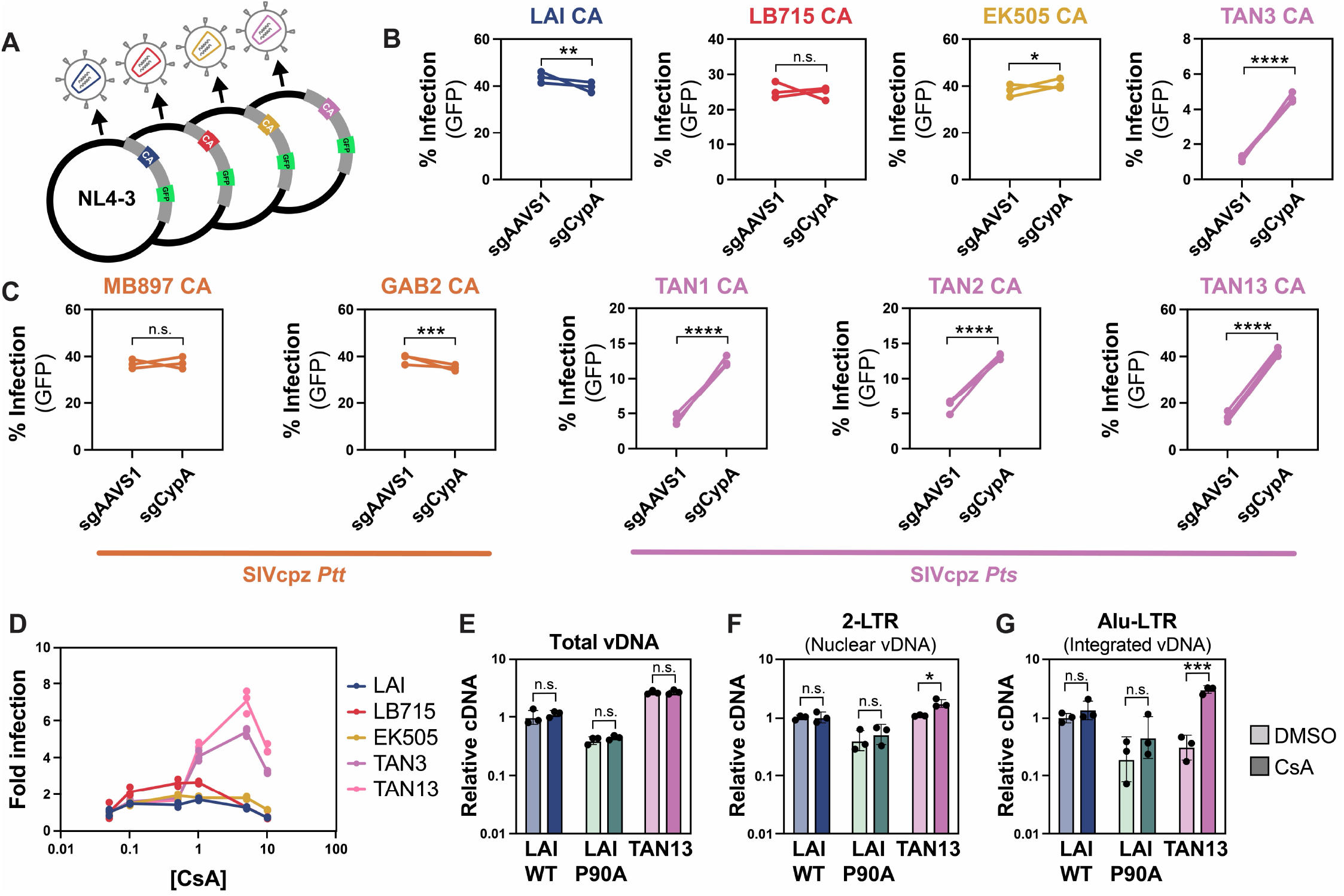
SIVcpz*Pts* capsids are restricted by CypA before integration. **(A)** Schematic of the capsid (CA) dropout HIV-1 NL4-3 ΔEnv, ΔNef GFP reporter vector system. (**B** and **C**) Infection levels of input-normalized CA reporter viruses as measured by GFP fluorescence in control *AAVS1* or CypA edited THP-1 cells. Infection with capsids from screened viruses in **B**. Additional SIVcpz*Ptt* and SIVcpz*Pts* capsids in **C. (D)** Relative infection of input-normalized CA chimera viruses in response to increasing doses of CypA inhibitor cyclosporin (CsA). Fold infection was determined by normalizing the percent infection (GFP flow cytometry) of each treatment group to the lowest tested dose of CsA (0.05 µM). (**E** to **G**) qPCR-based lentivirus lifecycle assays. Quantification of viral DNA products in DMSO or CsA (1 µM) treated cells 24 hours after infection with reporter virus encoding LAI CA (blue), LAI P90A CA (green) or TAN13 CA (purple) as measured by qPCR. Relative viral cDNA products were determined first by normalization to GAPDH and then normalized to DMSO-treated HIV-1 LAI infection. Infections were done in triplicate. **(E)** Total vDNA primers detect all viral DNA produced during reverse transcription. **(F)** 2-LTR primers detect viral DNA products that form only after entry into the nucleus. **(G)** Alu-LTR primers detect integrated proviral DNA by amplification of integrated viral DNA near commonly occurring genomic Alu elements. Data are presented as mean ± SD of triplicates from a single representative experiment. Experiments were performed three times with similar results. Statistical significance was determined by a two-tailed, unpaired Student’s t-test between CypA and *AAVS1* edited cells (**B** and **C**) or between DMSO and CsA treated cells (**E** to **G**). *P<0.05; **P<0.01; ***P<0.001; ****P<0.0001; n.s.= not significant.

To ask how CypA inhibits SIVcpz*Pts* capsid we measured the accumulation of viral DNA products after infection (Fig. 2, E to G; fig. S7). In all cases, CsA treatment leads to no significant changes in viral DNA products for HIV-1 or the the CypA-binding deficient P90A capsid [33] (Fig. 2, E, F, and G). Completion of reverse transcription of Eastern chimpanzee SIVcpz*Pts* capsid-encoding viruses is not affected by CypA as total viral DNA does not change with or without CsA treatment (Fig. 2E). Similarly, the formation of 2-LTR products, which are dependent on nuclear localized DNA repair machinery, does not dramatically increase upon treatment with CsA for SIVcpz*Pts* capsid-encoding virus (Fig. 2F). However, treatment with CsA leads to a ~6-fold rescue of integrated viral DNA for the SIVcpz*Pts* TAN13 capsid-encoding virus (Fig. 2G). Therefore, the restriction of SIVcpz*Pts* capsids by CypA occurs post-nuclear entry and prior to integration of the viral DNA.

### A novel capsid-CypA interaction defines sensitivity of SIVcpz*Pts* to CypA restriction

CypA is a prolyl isomerase and binds to lentiviral capsids through its active site [34]. The ‘CypA binding loop’, a region encoded between helices 4 and 5 of the HIV-1 capsid, mediates this interaction [35]. Sites Gly89 and Pro90 in capsid, which are required for CypA binding to HIV-1 capsid [33], are conserved between SIVcpz and HIV-1 although the surrounding capsid sequence shows some variation (Fig. 3A). To ask if this loop is required for CypA restriction of SIVcpz*Pts* capsid, we replaced the HIV-1 LAI CypA binding loop with the SIVcpz*Pts* CypA binding loop (teal in Fig. 3, A and B). We find that the SIVcpz*Pts* CypA loop does not confer restriction to HIV-1 and CypA knockout does not confer any rescue (Fig. 3C). Interestingly, SIVcpz*Pts* capsids encode a 3-amino acid insertion in the disordered loop between helices 6 and 7 relative to both SIVcpz*Ptt* and HIV-1 (red in Fig. 3, D and E). This 6,7 loop forms non-covalent interactions with the CypA binding loop in certain non-pandemic HIVs and SIVs [31]. Swapping the 6,7 loop from SIVcpz*Pts* capsid into HIV-1 LAI capsid is sufficient to confer restriction that is rescued upon CypA knockout (Fig. 3F) or after disrupting capsid-CypA interaction with CsA (Fig. 3G). Conversely, swapping the HIV-1 LAI capsid 6,7 loop sequence into SIVcpz*Pts* abrogates CypA-dependent restriction (Fig. 3H). Therefore, the SIVcpz*Pts* capsid 6,7 loop is both necessary and sufficient for CypA restriction. We also observe significant rescue of the SIVcpz*Pts* 6,7 loop swap virus in 2 donors in a primary CD4+ T cell model of HIV infection (fig. S8). To ask if the CypA restriction mediated by the SIVcpz*Pts* capsid 6,7 loop is dependent on CypA binding at its canonical binding loop, we introduced Ala at residue 90 in the SIVcpz*Pts* 6,7 loop swap virus. This mutation fully rescues the CypA restriction (Fig. 3I). In sum, CypA restriction is dependent on both the SIVcpz*Pts* 6,7 loop and on interaction at the canonical capsid-CypA binding interface.

**Figure 3.**
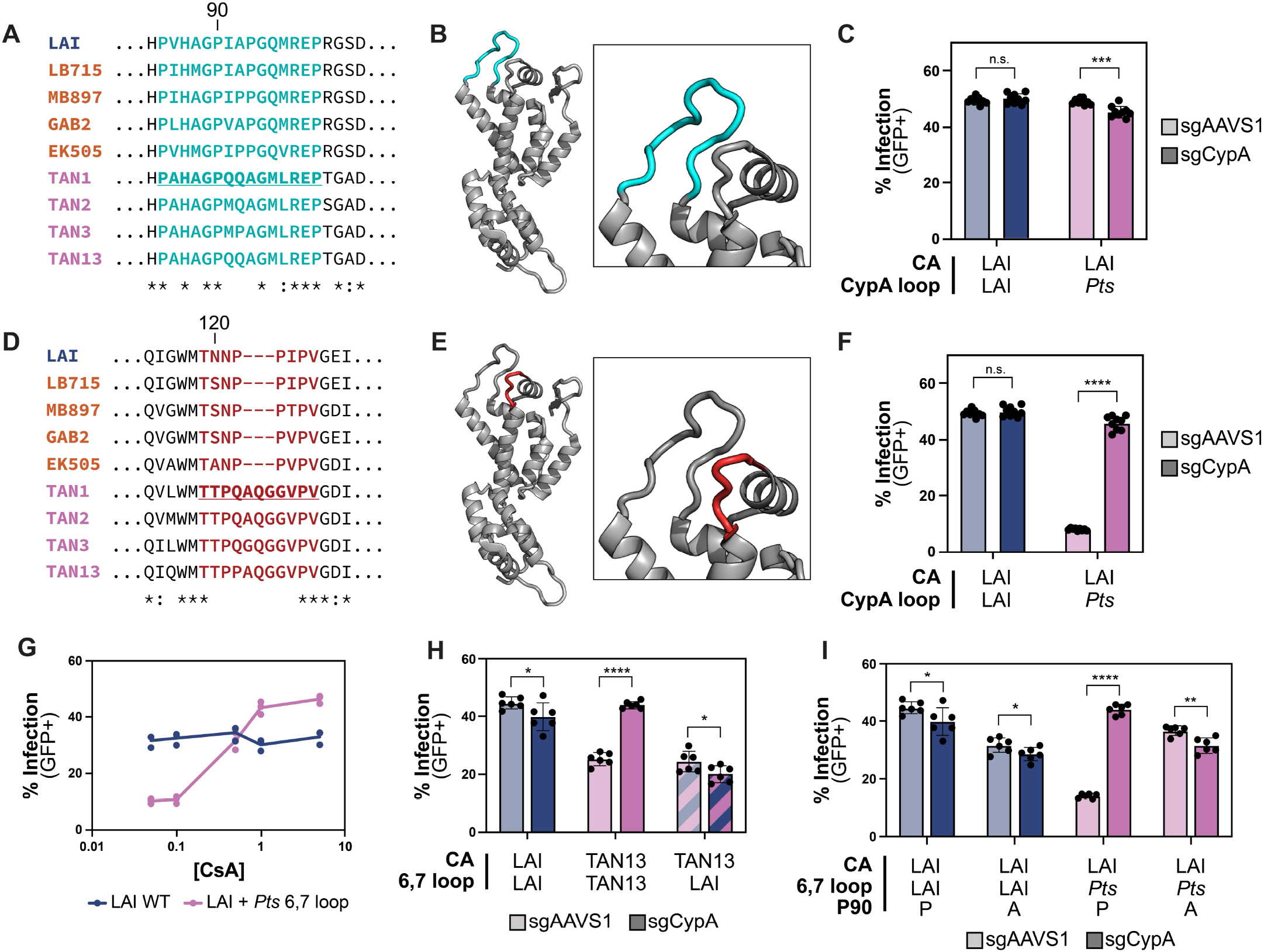
A unique insertion in the SIVcpz *Pts* capsid 6,7 loop underlies the CypA restriction phenotype. **(A)** Amino acid alignment of the CypA binding loop protein sequences (teal region) in SIVcpz and HIV-1 M capsids. The representative sequence cloned into the loop swap chimera is underlined. HIV-1 M in blue; Central chimpanzee SIVcpz*Ptt* in orange; Eastern chimpanzee SIVcpz*Pts* in purple. **(B)** HIV-1 M capsid monomer (PDB: 2M8N) with CypA binding loop highlighted in teal. **(C)** Infection levels of input-normalized wild-type LAI CA (blue) or LAI CA with the SIVcpz*Pts* CypA binding loop swap (purple) in *AAVS1* control or CypA edited THP-1 cells. **(D)** Alignment of the 6,7 loop protein sequences (red) in SIVcpz and HIV-1 M capsids. The representative SIVcpz*Pts* capsid sequence cloned into the loop swap chimera is underlined. **(E)** HIV-1 M capsid monomer with the 6,7 loop highlighted in red. **(F)** Infection levels of input-normalized LAI CA (blue) or LAI CA with the SIVcpz*Pts* 6,7 loop swap (purple) in *AAVS1* control or CypA edited THP-1 cells. **(G)** Infection levels of input-normalized LAI CA (blue) or LAI CA with the SIVcpz*Pts* 6,7 loop swap (purple) with increasing doses of CsA in THP-1 cells. **(H)** Infection levels of input-normalized HIV-1 LAI CA (blue), SIVcpz*Pts* TAN13 CA (purple) or TAN13 CA with the LAI 6,7 loop swap (blue and purple striped) in *AAVS1* control or CypA edited THP-1 cells. **(I)** Infection levels of input-normalized HIV-1 LAI CA (blue) or HIV-1 LAI CA with the SIVcpz*Pts* 6,7 loop swap (purple) with or without the CypA binding loop P90A capsid mutation in *AAVS1* control or CypA edited THP-1 cells. Data are presented as mean ± SD of triplicates from a single representative experiment. Experiments were performed three times with similar results. Statistical significance was determined by a two-tailed, unpaired Student’s t-test between CypA and *AAVS1* edited cells (**C, F, H** and **I**). *P<0.05; **P<0.01; ***P<0.001; ****P<0.0001; n.s.= not significant.

Next, we solved the x-ray crystal structures of HIV-1 LAI or HIV-1 SIVcpz*Pts* 6,7 loop swap chimeras in complex with CypA (Fig. 4, A to C; CypA in gray; fig. S9; Table S1). Co-crystallization shows two distinct capsid-CypA interface conformations for both the HIV-1 LAI and SIVcpz*Pts* 6,7 loop swap capsids (Fig. 4A). For both capsids, one interface shows no direct interaction between CypA and the 6,7 loop, consistent with previous structural data for HIV-1 capsid [35] (fig. S9, A and B). However, both capsids show another conformation with direct interaction between CypA and the 6,7 loop (Fig. 4, B and C). For HIV-1 LAI, the interaction between the 6,7 loop and CypA is mediated by Asn121 (Fig. 4B). In contrast to HIV-1 LAI, SIVcpz*Pts* 6,7 loop shows more substantial contacts with CypA including through residue Gln124, which together with the loop extension forms a unique interaction with Arg148 in CypA (Fig. 4C). Gln124 is not encoded in the 6,7 loop of either pandemic HIV-1 M or zoonotic SIVcpz*Ptt* capsid as this is within the unique insertion found only in SIVcpz*Pts* capsids (see Fig. 3D and fig. S12). Therefore, the SIVcpz*Pts* 6,7 loop sequence creates a unique interaction between capsid and CypA. To ask how this capsid-CypA interface modulates restriction, we introduced point mutations at residue 124 in the SIVcpz*Pts* 6,7 loop swap capsid and infected wild-type or CypA knockout THP-1 cells (fig. S10). Mutation of Q124 to glutamic acid (E) prevents restriction by CypA (Fig. 4, D and E). Therefore, residue 124 in the SIVcpz*Pts* 6,7 loop forms a unique interaction with CypA and determines CypA restriction.

**Figure 4.**
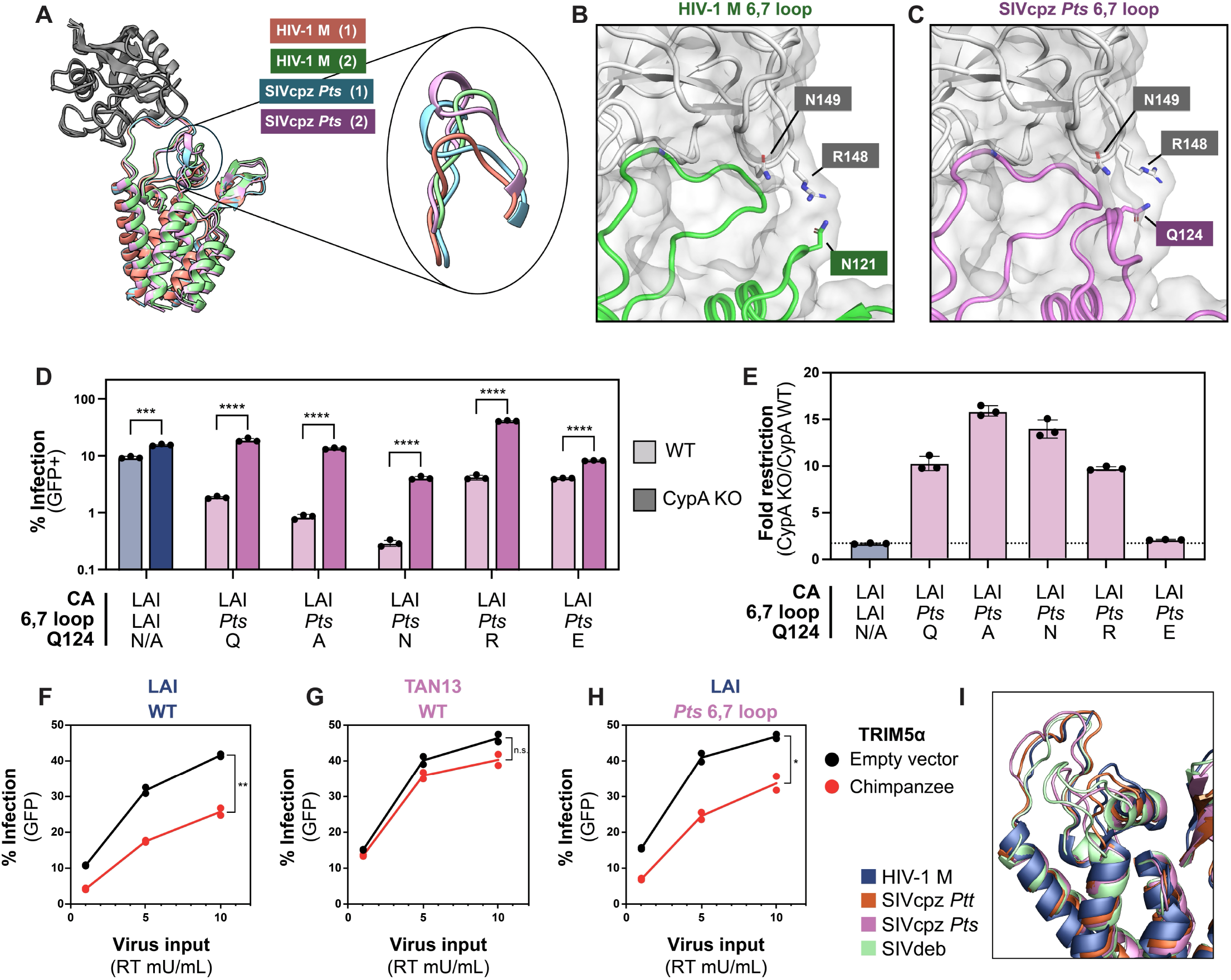
The Eastern chimpanzee SIVcpz*Pts* capsid forms a unique interface with CypA and evades chimpanzee TRIM5α restriction. (**A** to **C**) X-ray crystal structures of the N-terminal domain of HIV-1 LAI CA **(B)** and HIV-1 LAI CA with the SIVcpz*Pts* 6,7 loop swap **(C)** (all structures overlaid in **A**). (**B** and **C**) Close-up views of the interface between capsid and CypA, highlighting residues at the interaction site (R148 and N149 in CypA; N121 and Q124 in HIV-1 and SIVcpz*Pts* 6,7 loop capsid respectively). **(D)** Infection levels of input-normalized HIV-1 LAI CA (blue) or LAI CA chimera with the SIVcpz*Pts* 6,7 loop swap (purple) with the indicated single amino acid substitutions at position 124 in capsid (Q, wild-type; A, Alanine; N, Asparagine; R, Arginine; E, Glutamic acid) in wild-type or a CypA knockout THP-1 clonal cell line as detected by GFP flow cytometry. This experiment was performed once in clonal CypA knockout cells; an additional experiment using CsA showed similar results. **(E)** Relative restriction by CypA of capsid mutant viruses calculated as the ratio of percent infection in CypA knockout cells relative to wild-type cells (CypA KO / CypA WT). (**F** to **H**) Infection levels of reporter viruses encoding different capsids in human TRIM5α knockout THP-1 cells transduced with an empty vector (black) or chimpanzee TRIM5α overexpressing vector (red). HIV-1 LAI capsid in **F**. TAN13 capsid in **G**. HIV-1 LAI with the SIVcpz *Pts* 6,7 loop swap in **H. (I)** AlphaFold predicted structures of the capsid monomers of SIVcpz*Ptt* (orange), SIVcpz*Pts* (purple) and SIVdeb (green) overlaid with the X-ray crystal structure of HIV-1 M (PDB: 2M8N; blue). Data presented are replicates from a single representative experiment. Experiments were performed three times with similar results unless otherwise indicated. Statistical significance was determined by a two-tailed, unpaired Student’s t-test between CypA knockout or wild-type cells **(D)** or between empty vector and chimpanzee TRIM5α overexpressing cells at the maximum virus input concentration (10 mU/mL) **(H-J)**. *P<0.05; **P<0.01; ***P<0.001; ****P<0.0001; n.s.= not significant.

### Evolution of CypA-restricted SIV: tradeoffs and origins

To understand how the CypA-restricted capsid sequence evolved in Eastern chimpanzees, we looked at other capsidhost factor interactions that may have influenced its evolution. We first analyzed variation in CypA sequence within a new dataset across chimpanzee subspecies [36]. We found that CypA does not vary in amino acid sequence in the *P. t. troglodytes* chimpanzee reference genome as compared to humans. CypA gene sequences in Eastern and Central chimpanzee populations show low-frequency polymorphisms exclusively in noncoding regions that do not alter the CypA amino acid sequence (Table S2). Therefore, species specific differences in CypA do not explain the restriction of SIVcpz*Pts* capsids.

We next sought to determine if another host factor may have influenced the evolution of SIVcpz capsids. A major role for CypA in HIV-1 infection of human cells is to protect the HIV capsid from restriction by the host antiviral protein TRIM5ɑ that binds and restricts capsids [20,32,37]. In contrast to the lack of coding variation in CypA, TRIM5ɑ is a rapidly-evolving gene that varies in coding sequence between humans and chimpanzees and shows allelic variation within chimpanzee populations [38] (fig. S11, A and B; Table S2). To test the hypothesis that CypA-restricted SIVcpz*Pts* capsids may have been selected for by a restrictive chimpanzee TRIM5ɑ allele, we infected cells overexpressing chimpanzee TRIM5ɑ. The SIVcpz*Pts* TAN13 capsid virus is less sensitive to chimpanzee TRIM5ɑ restriction as compared to the HIV-1 M capsid (Fig. 4, F and G). Interestingly, the resistance to TRIM5ɑ restriction does not map exclusively to the 6,7 loop, as the 6,7 loop swap virus is as sensitive to TRIM5ɑ as the HIV-1 LAI capsid (Fig. 4H). Therefore, the 6,7 loop alone determines sensitivity to CypA restriction, but is not sufficient for resistance to TRIM5ɑ restriction.

To understand the evolutionary origins of the 6,7 loop insertion in Eastern chimpanzee SIV capsid, we aligned SIV capsid sequences and find that the extended 6,7 loop that confers CypA restriction in SIVcpz*Pts* is similarly extended in De Brazza’s monkey SIV capsid (SIVdeb) (fig. S12, A and B). Alphafold-predicted structures of SIVcpz*Ptt* (LB715 in orange), SIVcpz*Pts* (TAN3 in purple) and SIVdeb (green) capsids overlaid with the crystal structure of HIV-1 M (blue; PDB: 2MN8) show that the SIVdeb capsid structurally resembles that of SIVcpz*Pts* including a similarly extended 6,7 loop sequence (Fig. 4I) [39]. Acquisition of new sequence in lentiviruses is well known to occur through recombination following super-infection [14,40]. Therefore, the shared extended 6,7 loop conformations of SIVcpz*Pts* and SIVdeb may be an example of either shared ancestry due to a recombination event or of convergent evolution in the face of selection from CypA and TRIM5ɑ during engagement with the lentiviral capsid.

## Discussion

Here we show that CypA, a host factor that facilitates HIV infection, is a barrier to the replication of SIVcpz viruses en-demic to Eastern chimpanzees in human cells. This finding represents the first potential molecular explanation as to why SIVcpz*Ptt* and not SIVcpz*Pts* has crossed the species barrier, giving rise to HIV-1. Our study adds further support to the idea that the interaction with CypA is an important point of selection and adaptation in lentiviral capsids. Lentiviral capsid co-evolution with host CypA and related interactors is complex. Multiple lineages of primates have independently co-opted the capsid binding activity of CypA through evolved, novel TRIM-CypA gene fusions [41]. The dynamic nature of the capsid/CypA coevolution is also apparent in the recurrent gain and loss of binding to CypA in lentiviruses [42]. Interestingly, many primate lentiviruses also rely on the cyclophilin binding activities of capsid to enter the nucleus using RanBP2, a nuclear pore protein with a cytoplasmic cyclophilin domain [30]. Our observation that CypA restricts SIVcpz*Pts* after nuclear entry (Fig. 2G) offers a tempting hypothesis that CypA could guard against nuclear entry of certain lentiviruses. As such, the toggling across evolution of dependency and restriction of capsids by cyclophilin binding would be an expected outcome.

One potential evolutionary explanation for the emergence of the CypA-restricted capsid in Eastern chimpanzees is selection on this capsid for resistance to restriction by chimpanzee TRIM5ɑ. Like many canonical restriction factors, the characteristic rapid evolution and selection on TRIM5ɑ has resulted in functionally important variation between even very closely related primates [38]. Our results suggest an evolutionary tradeoff in which SIVcpz*Pts* capsids are sensitive to inhibition by CypA but less restricted by chimpanzee TRIM5ɑ. In this model, chimpanzee TRIM5ɑ may have selected for SIVcpz*Pts* capsids with evolved TRIM5ɑ resistance at the cost of CypA sensitivity. In contrast, SIV from Central chimpanzees appears to have maintained the co-option of CypA as protection from TRIM5ɑ restriction, which in turn makes these capsids more amenable to infecting humans. Our results highlight the complex nature of multiple host factors driving competing evolutionary trajectories on the same capsid sequence. In sum, CypA presents a barrier to infection in human cells that distinguishes nonzoonotic and zoonotic chimpanzee lentiviruses.

When a host is contemporaneously infected with more than one lentivirus, recombination serves as a source for rapid adaptation to novel host factors. These recombination events are known to have played an important role in the evolution of pandemic HIV-1 [40]. De Brazza monkey, genus *Cercopithecus*, harbors diverse SIVs and serves as prey for chimpanzees [43]. We speculate that the 6,7 loop extensions shared between Eastern chimpanzee SIV and De Brazza monkey viral capsids could be the result of an ancestral recombination event between these viruses. The evolution of this CypA-restricted SIVcpz*Pts* capsid may explain the lack of efficient transmission of this clade of SIVcpz to humans. The discovery of host CypA as a restriction factor of SIVcpz*Pts* capsids presents a novel mode of host defense against lentiviruses with important consequences for zoonotic spillover potential.

## Acknowledgments

We thank Jeannette Tenthorey for providing the clonal TRIM5α knockout THP-1 cells as well as the TRIM5α overexpression vector system. We thank Beatrice Hahn and Frederic Bibollet-Ruche for the SIV infectious molecular clones and for helpful discussion. We thank Russell Vance and Jeanette Tenthorey for helpful feedback on the manuscript. We thank Ohainle Lab members including Bridgett Rios, Isaiah Grant, Leticia Pereira and Nicholas Goodin as well as the Berkeley MCB community for helpful discussions. The content is solely the responsibility of the authors and does not necessarily represent the official views of the National Institutes of Health.

## Author Contributions

Conceptualization: MJY, MO; Methodology: MJY, FC, CCCG, JAL, FGW, AA, JLR, MS, PHS, OP, MO; Investigation: MJY, FC, JAL, JLR, MS, OP, PHS, MO; Visualization: MJY, AA, MAN, FF, JAL, MO; Funding acquisition: MAN, OP, PHS, MO; Project administration: MO; Supervision: OP, PHS, MO; Writing – original draft: MJY, MO; Writing – review & editing: MJY, FC, CCCG, JAL, AA, JLR, MS, MAN, FF, PHS, OP, MO.

## Data Availability

The atomic models generated in this study have been deposited in the Protein Data Bank (PDB): HIV-1 LAI capsid in complex with CypA (PDB PDB_000038IH) and HIV-1 LAI capsid with the SIVcpz*Pts* 6,7 loop swap in complex with CypA (PDB PDB_000038IG). All raw and minimally-processed HIV-CRISPR screen data are available through the NCBI Gene Expression Omnibus (GEO): GSE343775. Processed HIV-CRISPR screen data is available as Supplementary Material (Data S1-S5) and can be viewed at www.crisprvirus.org. Code used for *Pan troglodytes* genomic analysis can be found in the following GitHub repository: https://github.com/sudmantlab/panpan_diversity_project. The vcf files used for SNP detection in candidate genes of interest are archived in zenodo (PENDING). All raw long-read sequencing data are deposited in NCBI under accession number (PENDING). Genome assemblies are deposited under (PENDING). Chimpanzee short-read sequencing data includes publicly available data from NCBI bioprojects PRJEB15086 and PRJNA189439. All data analyses were performed using published computational pipelines and available python packages as described in the Materials and Methods. Custom code for NTC reconstruction and post-MAGeCK processing is available on Github at https://github.com/ohainlelab/crispr-count-remapper.

## Materials & Methods

### Phylogenetic trees and alignments

For generation of the SIVcpz cladogram, the following HIV and SIVcpz sequences were obtained from the Los Alamos National Laboratory (LANL) HIV Database: HIV-1 M LAI (HIVBRUCG), SIVcpz LB715 (JX178450), SIVcpz EK505 (DQ373065), SIVcpz TAN3.1 (DQ374658), SIVcpz MB897 (EF535994), SIVcpz GAB2 (AF382828), SIVcpz TAN1 (AF447763), SIVcpz TAN2.69 (DQ374657), SIVcpz TAN13 (JQ768416). Multiple sequence alignment (MSA) of full HIV and SIVcpz nucleotide genomes was generated using MAFFT in PhyML via the NGPhylogeny.fr web server with default settings. A maximum-likelihood tree was then generated using 100 bootstrap replicates and visualized in FigTree (v1.4.4). For HIV and SIV capsid protein alignments, nucleotide genomes of HIV and SIV were obtained from Bell & Bedford 2017 [1] and the LANL HIV database. Regions of capsid were determined based on homology and then translated to amino acid sequence. Capsid protein MSAs were then generated using Clustal Omega (v1.2.4) via the EMBL-EBI web server with default settings.

### Cell culture

Human THP-1, K-562, Jurkat, HuT-78, SupT1, CCRF-CEM cells and 293T cells were obtained from ATCC and cultured in RPMI (Gibco) with 10% FBS, Pen/Strep, 10 mM HEPES, 0.11 g/L sodium pyruvate, 4.5 g/L D-Glucose and Glutamax. HEK293T cells were cultured in DMEM (Gibco) with 10% FBS and Pen/Strep. Limiting dilution in round bottom 96-well plates was used for generation of THP-1 clonal cell lines.

### Plasmids

The HIV-CRISPR vector (NIH HIV Reagent Program; ARP-13567) was previously described [2,3]. pMD2.G and psPAX2 plasmids were a gift from Didier Trono (Addgene; 12259/12260). lentiCRISPRv2 was a gift from Feng Zhang (Addgene; 52961). Cloning of sgRNAs targeting genes of interest was performed using BsmBI restriction cloning as previously described [2]. HIV-1 LAI and HIV-1 LAI Δenv are previously described [2]. SIVcpz (LB715, MB897, GAB2, EK505, TAN1.910, TAN2.69, TAN3.1, TAN13) infectious molecular clones (IMCs) were a gift from Beatrice Hahn. For generation of Δenv SIVcpz viruses (LB715, EK505, TAN3), a 2 base pair deletion was introduced into the envelope coding sequence by PCR creating a frameshift mutation and cloned into the IMCs using BstBI and SbfI (LB715), SbfI and BamHI (EK505) and AgeI and NdeI (TAN3). Sequences are in Table S3. The NL4-3 GFP Δenv molecular clone was a gift from Jeremy Luban [4]. To create pNL-CAgg Δenv GFP, P_lac_-eGFP was amplified from pUCBB-eGFP (a gift from Russell Vance) with external BsmBI restriction sites by PCR and assembled in place of capsid in the NL43 GFP Δenv molecular clone by Gibson assembly. For CypA binding loop swap, 6,7 loop swap, or mutant capsid sequences, capsid sequences were synthesized by Twist Bioscience to include BsmBI cut sites. Capsid sequences from HIV or SIVcpz were amplified by PCR to incorporate BsmBI cutsites for cloning into pNL-CAgg-Δenv-GFP. The lentiviral transfer vector for overexpressing human TRIM5ɑ under the EF1ɑ full promoter (pHIV-hsTRIM5-HA-P2A-mCherry-IRES-puro) or equivalent empty vector (pHIV-empty-P2A-mCherry-IRES-puro) were gifts from Jeannette Tenthorey. These vectors were created from pHIV-ZsGreen (Addgene; 18121) using the XbaI and ClaI cut sites. The CSII-IDR2-chimpTRIM5-FOS vector encoding chimpanzee TRIM5ɑ was a gift from Wesley Sundquist (Addgene; 79065). The chimpanzee TRIM5ɑ sequence in CSII-IDR2-chimpTRIM5-FOS was amplified by PCR to add an HA-tag and NotI and NheI cut sites and ligated into pHIV-empty-P2A-mCherry-IRES-puro to create pHIV-ptTRIM5-P2A-mCherry-IRES-puro.

### Lentivirus production

For all transfections, 293T cells were seeded at 5 x 10^5^ cells/well in 2 mL in 6-well plates one day before transfection. Transfections were performed using 3 µL of TransIT-LT1 reagent (Mirus Bio; MIR2304) per µg of DNA. lentiCRISPRv2 and pHIV transfections were performed using 667 ng of transfer plasmid, 500 ng psPAX2 and 333 ng of pMD2.G. For production of all full-length native HIV envelope HIV or SIVcpz viruses, 293T cells were transfected with 1 µg of virus plasmid. For production of Δenv HIV and SIVcpz virus pseudotyped with VSV-G, 293T cells were transfected with 1 µg of virus plasmid and 333 ng of pMD2.G. For engineered Δenv molecular clones, the lack of native envelope expression was sequence confirmed and experimentally confirmed by testing infection with virus generated from transfection without pMD2.G. One day after transfection, the supernatant in each well was replaced with 1.5 mL complete DMEM. Two-days after transfection, the cell supernatant was collected and clarified by vacuum filtration through a 0.45 µm filter. Virus was then aliquoted and stored at −80ºC. Generation of lentiviral libraries for HIV-CRISPR screening was performed as previously described [2].

### Virus Infections

For infection and transduction in cell lines, cells were seeded at 1-2 x 10^5^ cells/well in 96-well flat bottom plates with DEAE-Dextran ([20ug/mL] Final) in RPMI. Virus or lentivirus was added and cultures were spinoculated at 1,000 x g for 20 minutes and incubated at 37ºC. For cells transduced with lentiviral particles (lentiCRISPRv2 or pHIV overexpression vectors) media was replaced the following day. Antibiotic selection was then initiated 1-3 days after transduction using 0.5-1 µg/mL puromycin. Transduction with TRIM5α overexpression vectors was validated by mCherry expression via flow cytometry. For infection with HIV or SIVcpz, cells were infected with HIV, SIVcpz or capsid chimeras normalized by reverse transcriptase activity (SG-PERT) where indicated. Infected cells were immediately transferred to V-bottom 96-well plates following spinoculation and spun at 1,000 x g for 3 minutes. The infected cell pellets were then resuspended in fresh RPMI and transferred to a 96-well flat bottom plate and incubated at 37ºC. For experiments using cyclosporin (CsA; Sigma-Aldrich SML1018), cells were resuspended in CsA media on resuspension following centrifugation in the V-bottom plate. Cells were fixed and analyzed for GFP or intracellular p24 expression by flow cytometry.

### Flow cytometry

For detection of GFP-positive cells, cells were pelleted in V-bottom 96-well plates at 1000 x g for 3 minutes and cell pellets were fixed by resuspension in 40 µL CytoFix at 4% PFA (BD Cytofix Fixation Buffer; 554655). Cells were incubated at room temperature for 10 minutes before diluting to 1% PFA with PBS. For intracellular p24 staining, cells were permeabilized with CytoPerm (BD CytoFix/CytoPerm Fixation/Permeabilization Kit; 554714) and stained with a 1:300 dilution of KC57-RD1 antibody (Beckman Coulter; 6604667). Cells were read on a Beckman CytoFlex Flow Cytometer and analyzed for GFP or intracellular p24 expression in FlowJo v10.10.0.

### SG-PERT Viral release assay

Viral release was determined by detection of reverse transcriptase activity in viral supernatants using the SG-PERT assay [5]. In brief, reverse transcriptase activity was determined by qPCR using the QuantStudio 6 Pro (Applied Biosystems). Reverse transcriptase (RT) units were calculated using a standard curve of HIV-1 LAI standard virus with calculated RT activity units calculated from purified, recombinant RT.

### HIV-CRISPR screening

HIV-CRISPR sgRNA library assembly and screening were performed as described previously [2,3]. For genome-wide screens, the human Brunello CRISPR knockout library (Addgene; 73179) was cloned into the HIV-CRISPR vector as described previously [2]. To generate the ZIRA sublibrary, the 300 highest-ranked putative restriction factors and 300 highest-ranked putative dependency factors from each of four genome-wide HIV-CRISPR screens performed with LAI, LB715, EK505, and TAN3 were pooled and deduplicated, yielding 1,477 genes. Ten sgRNAs per gene were designed with CRISPick (53, 54). The resulting library contained 16,000 sgRNAs: 14,770 gene-targeting sgRNAs and 1,230 non-targeting control (NTC) sgRNAs. ZIRA library is in Data S3. The oligonucleotide pool was synthesized by Twist Bioscience and cloned into the HIV-CRISPR vector as described previously [2]. Brunello or ZIRA THP-1 cell libraries were infected with each virus to achieve 5-40% infection. All screens were performed in duplicate. Screening data were analyzed with MAGeCK v0.5.9.5 [6]. FASTQ reads were processed with cutadapt v2.6 [7] to remove the 5′ constant sequence CTTGTGGAAAGGACGAAACACCG and retain the 20-nucleotide sgRNA sequence. Processed reads were mapped to the corresponding library, and sgRNA abundances were quantified with ‘mageck count’. Details on data processing available on GitHub: https://github.com/ohainlelab/crispr-count-remapper. Briefly, NTC sgRNAs were used to generate equal number of ‘synNTC’ genes as genes in the library with equivalent NTC sgRNAs per ‘synNTC’ gene as other genes in the library.

Read counts were normalized using MAGeCK’s default median normalization. For each screen, replicate viral RNA (vRNA) samples were designated as the treatment group and compared with the corresponding genomic DNA (gDNA) reference samples using ‘mageck test’. MAGeCK ranked sgRNAs by their differential representation between vRNA and gDNA and used robust rank aggregation to calculate gene-level positive- and negative-selection statistics. Negative selection identified sgRNAs depleted from vRNA relative to gDNA, whereas positive selection identified sgRNAs enriched in vRNA. For visualization and candidate ranking, MAGeCK RRA scores were converted to −log10 values, with negative and positive scores indicating depletion and enrichment, respectively. NTCs were retained as empirical background references in screen visualizations but were not used for count normalization. Whole genome and ZIRA sgRNA scores are in Data S2 and S5. Gene scores are in Data S1 and S4.

### CRISPRvirus Data Commons

To make the large dataset generated in this study publicly accessible, and to facilitate visualization, interactive exploration, and direct comparison with functional genomics screening data in a broad range of cell types generated by many research groups, we developed CRISPRvirus (www.crisprvirus.org). CRISPRvirus was developed based on infrastructure from CRISPRbrain (www.crisprbrain.org) [8] and CRISPRlipid (www.crisprlipid.org) [9]. Data included in CRISPRvirus was analyzed using custom MAGeCK analysis pipelines as described [8,9].

### Pooled and clonal gene knockout

For nucleofection, 2 x 10^5^ THP-1 cells were electroporated with sgRNAs (EditCo Gene Knockout Kits) targeting *BST2* (Tetherin), *PPIA* (CypA) or *TRIM5* and purified recombinant Cas9 (UC Berkeley QB3 MacroLab) using the SG Cell Line 96-well Nucleofector Kit (Lonza Bioscience; V4SC-3096). Cells were immediately diluted in fresh media and allowed to recover for several days before generating single cell clones by limiting dilution. For pooled knockouts using lentiCRISPRv2, knockouts were generated as previously described [2]. Briefly, sgRNA oligos targeting genes of interest were ordered from Integrated DNA Technologies and annealed and ligated into the lentiCRISPRv2 lentiviral vector encoding puromycin resistance. Oligo sequences are in Table S3. THP-1 cells were then transduced and selected in 1 µg/mL puromycin for greater than 1 week. For both nucleofected and lentiCRISPRv2 edited cells, knockout was validated by western blot.

### Western blot

2-3 x 10^6^ cells were pelleted by centrifugation at 1000 x *g* for 3 minutes at 4ºC and lysed with NP-40 buffer (Fischer Scientific; AAJ60766AP) treated with Protease Inhibitor Cocktail (Roche; 11836170001). Lysates were clarified by centrifugation and boiled in a 6X SDS-PAGE loading buffer (G Biosciences; 10153-804) with 2-mercaptoethanol and boiled at 95ºC for 7 minutes. Proteins were resolved using NuPAGE 4-12% Bis-Tris mini gels (Invitrogen; NP0322BOX). Samples were run at 100V for 1-2 hours in 1X NuPAGE MES SDS Running Buffer (Fisher Scientific; NP0002) and transferred to a nitrocellulose membrane. Membranes were blocked in 2% milk and 2% BSA Tris-buffered saline with Tween 20 (TBST) at room temperature for 1 hour. Membranes were then incubated with primary antibody for 2 hours at room temperature or overnight at 4ºC. Membranes were then incubated with a secondary antibody for 1 hour at room temperature before imaging on LiCor Odyssey CLx. Protein expression was quantified relative to GAPDH in ImageJ 1.54g. For Tetherin western blotting, cells were treated with 1000 U/mL universal IFNα (PBL Assay Science; 11200) 24 hours before collection. Antibodies used: *BST2*/Tetherin at 1:8000 (Proteintech; 13560-1-AP), *PPIA*/CypA at 1:500 (CST; 2175S), ɑ-Actinin at 1:1000 (BioRad; VPA00889), GAPDH at 1:2000 (BioRad; AHP1628) and IRDye 800CW at 1:20,000 (LiCor; 926-32211).

### HIV lifecycle assays

HIV lifecycle assays were performed slightly modified from [10]. Briefly, 1 x 10^6^ cells per replicate were infected with DNAse (Promega; M6101) treated virus in 48-well plates and treated with DMSO or 1 µM CsA. Cells were washed twice with PBS and genomic DNA was extracted with the QIAamp DNA Blood Mini Kit (Qiagen; 51104) 24 hours after infection with RT-normalized HIV or SIV. Primer sets targeting reverse transcribed viral DNA (total viral DNA), circularized proviral DNA (2-LTR) or integrated proviral DNA (Alu-LTR) were used to amplify each product respectively (Table S3). Total viral DNA and 2-LTR products were detected with Takyon ROX SYBR Mastermix (Eurogentec; UF-RSMT-B0710) or iTaq Universal SYBR Green (BioRad; 1725121) with 2.5 uL of template DNA and 500 nM of each primer. GAPDH reactions for each sample were performed in parallel. Amplifcation was performed using the QuantStudio 6 Pro (Applied Biosystems) at 50ºC for 2 min, then 95ºC for 10 min, followed by 40 cycles of 95ºC for 15 sec and 62ºC for 1 min. Alu-LTR products were detected using a semi-nested PCR. Round one amplification was performed with Herculase (Agilent Technologies; 600677) with 200 ng of template DNA and 250 nM of each round one primer. Reactions were cycled in VeritiPro Thermal Cycler at 95ºC for 5 min, followed by 20 cycles of 95ºC for 30 sec, 52ºC for 30 sec, 72ºC for 4 min, and a final 72ºC for 10 min. PCR products were purified using AMPure beads (UC-Berkeley DNA sequencing facility). Round 1 PCR products were used in a second amplification with iTAQ Universal Probes Supermix (Biorad; 1725130) with 4 µL of round one product, with 300 nM of each primer and 100 nM of probe. Amplification was performed using the QuantStudio 6 Pro with the following cycling conditions: 95ºC for 3 min, followed by 40 cycles of 95ºC for 15 sec, then 60ºC for 90 sec. In parallel a reaction using the 20X GAPDH PrimeTime Probe Mix (IDT; Hs.PT.58.589810.g) with iTaq Universal Probes master mix and 4 uL of round one template was performed.

### Primary cell culture and infection

Peripheral blood mononuclear cells (PBMCs) were isolated from whole blood Leukopaks (Stanford Blood Center; B1005V00) from healthy human donors by density gradient centrifugation using Ficoll-Paque PLUS (Cytiva; 17144002). Primary human CD4+ T cells were isolated from PBMCs by negative selection using the EasySep Human CD4+ T Cell Isolation Kit (StemCell Technologies; 17952) according to the manufacturer’s protocol. Isolated CD4+ T cells were subsequently activated in complete RPMI-1640 supplemented with 100 U/mL IL-2 (Thermo Fisher Scientific #200-02-100UG) and 5 µg/mL anti-CD28 (Tonbo Biosciences; 40-0289) in 24-well plates pre-coated with 10 µg/mL anti-CD3 (Tonbo Biosciences # 40-0038). Cells were maintained in complete RPMI supplemented with 100 U/mL IL-2 for 8 days post-activation prior to infection. Primary human CD4+ T cells were infected on day 8 post-activation in 96-well plates with 50 RT mU/mL virus as described above and treated with DMSO or 1 µM CsA. Cells were fixed 2 days post-infection and analyzed for GFP expression by flow cytometry.

### X-ray crystal structures

HIV-1 M LAI capsid and the SIVcpz*Pts* 6,7 loop swap capsid chimera sequences (N-terminal domain only) were subcloned into pET11a, expressed, and purified using previously published methods [11,12]. CypA was expressed and purified as previously described [11]. Crystals were grown using the hanging drop vapor diffusion method by mixing the reservoir solution (100 mM Bicine pH 7, 1 M LiCl, and 20% Polyethylene Glycol 8000) with the protein solution (250 mM CypA and 250 mM CA in 10 mM Tris pH 8, 1 mM 2-mercaptoethanol). Crystals appeared within 24 hours and reached maximum growth after one week. Diffraction data were collected in-house using a Rigaku Synergy DW Diffractometer, scaled using XDS, and the phase problem was solved by using molecular replacement (PDB ID 1AK4). Model building was done iteratively using COOT [13] and Phenix [14]. X-ray data and structure determination statistics are in Table S1.

### Identification of SNVs in CypA and TRIM5

The *PPIA* (CypA) and *TRIM5* genomic sequences were analyzed in 20 *P. t. troglodytes* and 22 *P. t. schweinfurthii* chimpanzees using a vcf file of jointly called variants from novel long-read (n=24) and previously generated short-read (n=59) chimpanzee whole-genome sequencing data mapped to the T2T mPanTro3 chimpanzee reference (see [15] for method details on variant discovery). Data is in Table S2.

### AlphaFold predictions

The AlphaFold 3 Server [16] was used to predict the capsid structures of SIVcpz*Ptt* (LB715; JX178450), SIVcpz*Pts* (TAN3.1; DQ374658) or SIVdeb (04cmpf3061; FJ919724) using default parameters. Viral sequences were obtained from plasmid sequences or the Los Alamos National Laboratory HIV Database. Capsid sequences are in Table S3. Structural predictions were overlaid with HIV-1 M (PDB: 2MN8) in PyMOL (v3.1.6.1).

### Statistical Analysis

Statistical analysis was performed in GraphPad Prism 11. Statistical tests are indicated in the figure legends. Each data point is from an independent replicate from a representative experiment.

### Supplementary Materials

**Data S1**. Brunello MAGeCK gene score summary.

**Data S2**. Brunello MAGeCK sgRNA score summary.

**Data S3**. ZIRA sgRNA library.

**Data S4**. ZIRA MAGeCK gene score summary.

**Data S5**. ZIRA MAGeCK sgRNA score summary.

**Table S1**. X-ray crystal structure data.

**Table S2**. *PPIA* and *TRIM5* variation in *P. t. troglodtyes* and *P. t. schweinfurthii*.

**Table S3**. Relevant sequences used in study

