## Supplemental Figures for "CypA is a molecular barrier to HIV emergence from Eastern chimpanzees"

### Supplementary Figures

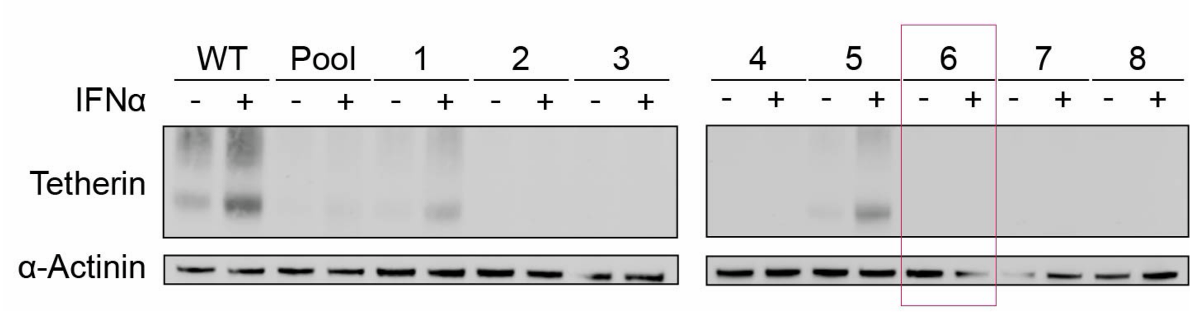


**Figure S1. Tetherin knockout THP-1 cell pools.** THP-1 cells electroporated with crRNPs targeting Tetherin (*BST2*) were single-cell cloned and clonal knockout of Tetherin in THP-1 cells validated by immunoblot. THP-1 cells were treated with or without universal IFNα for 24 hours before collection to stimulate Tetherin expression. Blots were probed for Tetherin and α-Actinin as a loading control. Clonal line used for subsequent experiments outlined in red.

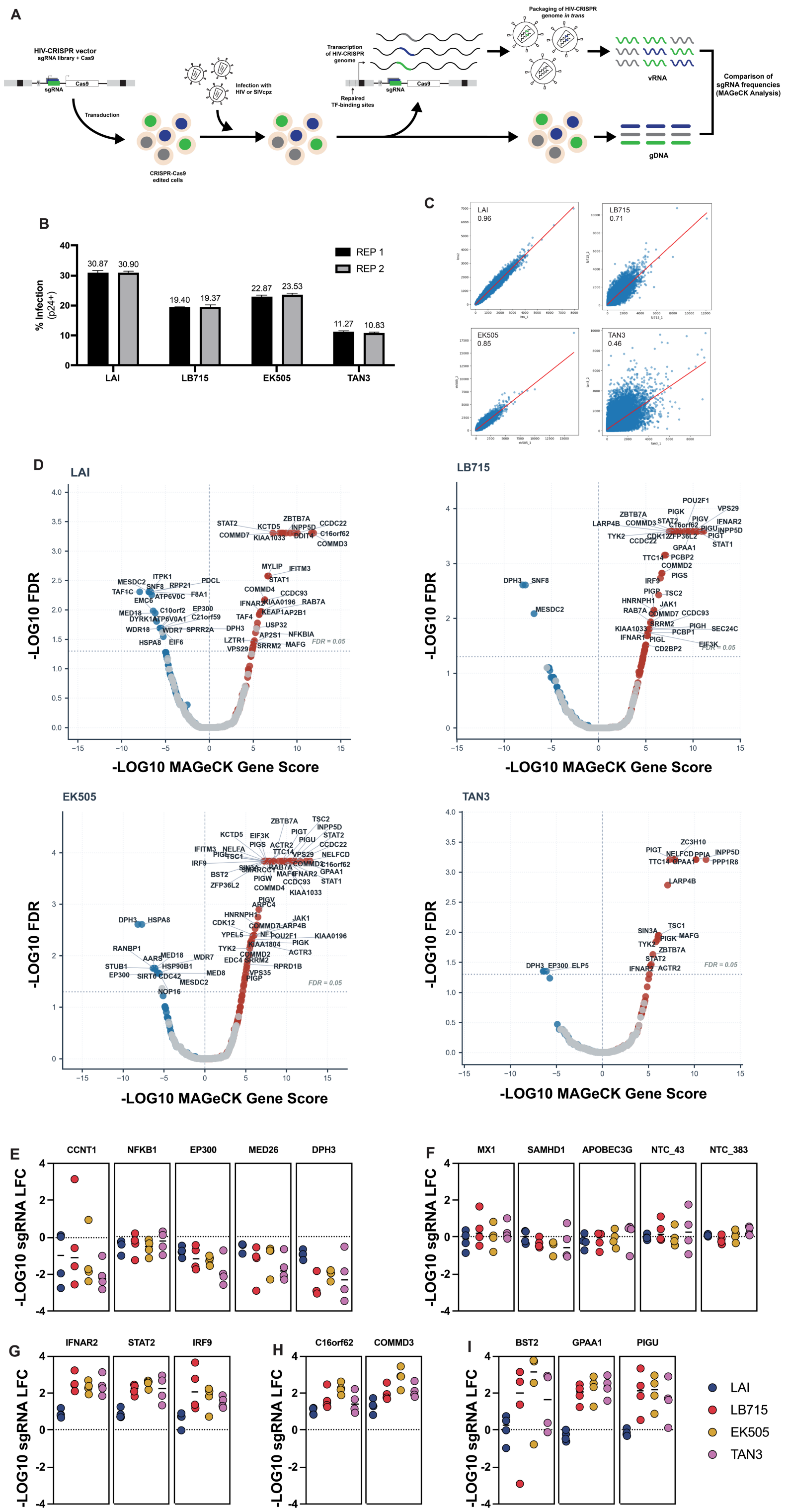

**Figure S2. Genome-wide HIV-CRISPR screens. (A)** Overview of HIV-CRISPR screening (*17*). (**B**) Infection levels as detected by intracellular p24 staining of genome-wide HIV-CRISPR screens across replicates. (**C**) Correlation of sgRNA counts between replicates for each genome-wide screen with indicated $R^2$ values. (**D**) Genome-wide HIV-CRISPR screen results with -log10 MAGeCK Gene Score (x-axis) plotted against -log10 FDR (y-axis). Genes where the -log10 MAGecK gene score > 0 are colored in red, < 0 colored blue, and NTCs are colored gray. Source data are in Data S1. (**E** to **I**) sgRNA -log10 fold change (LFC) scores derived using MAGeCK analysis for HIV-1 M (LAI in blue), SIVcpz*Ptt* (LB715 and EK505 in red and yellow) and SIVcpz*Pts* (TAN3) in purple. Source data are in Data S2. (**F**) Proviral host factors (CCNT1*, NFKB1*, EP300*, MED26*, DPH3). (**G**) Non-targeting controls ("NTC") and negative control genes (Mx1/MxA, SAMHD1*, APOBEC3G*). (**H**) Interferon Pathway (IFNAR2*, STAT2*, IRF9*). (**I**) Endosomal sorting (C16orf62, COMMD3). (**I**) Antiviral factors (Tetherin/*BST2**, GPAA1, PIGU). *Previously implicated in HIV or lentiviral replication.

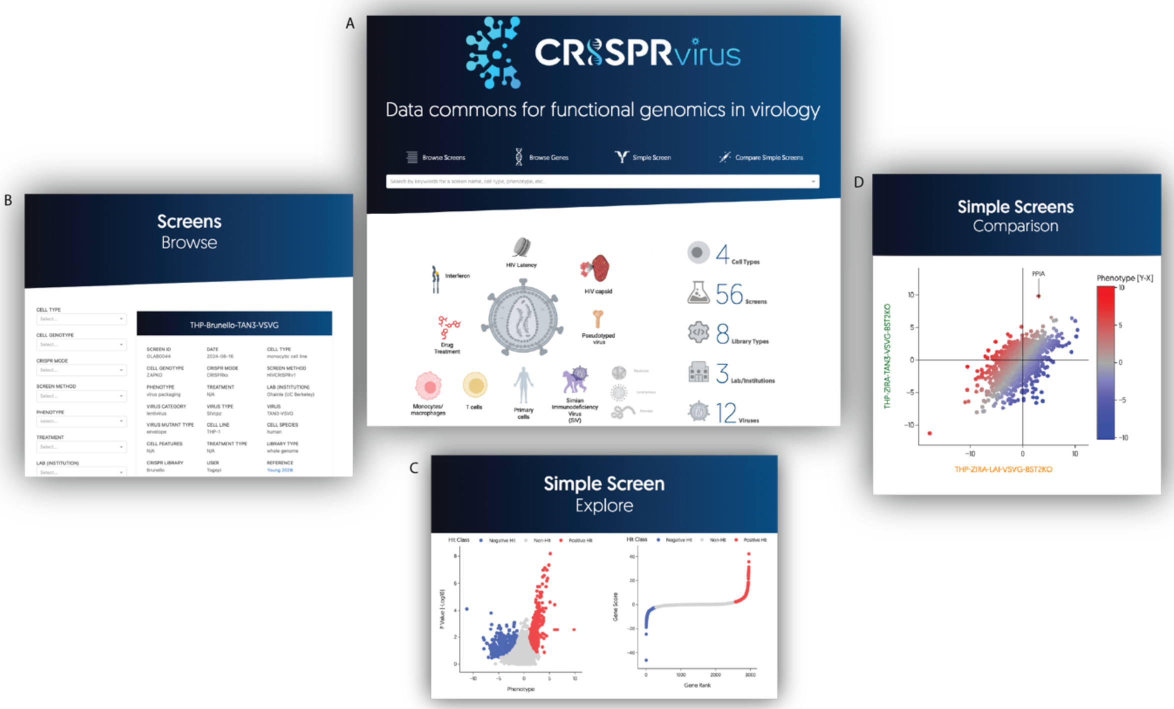


**Figure S3. CRISPRvirus: a data commons for CRISPR-based functional genomics screens in virology.** CRISPRvirus serves as a tool for rapid and easy comparison of functional genomic data in virology. **(A)** CRISPRvirus allows for interactive viewing and plotting of data from previous HIV-CRISPR screens along with the HIV-CRISPR screens from this study. (**B**) Screens can be filtered on a range of parameters including library, cell type, treatment, etc. (**C**) Screen data in CRISPRvirus can be easily visualized as scatterplots and plots and data tables can be downloaded. (**D**) Comparisons across screens are made easy as all data in the CRISPRvirus data commons are analyzed under the same parameters and with the same outputs. Current datasets available for exploration on CRISPRvirus include: screens in different cell systems including primary CD4+ T-cells (*17*, *21*); screens performed in HIV latency models including screens with latency reversal agents (LRA) (*44*–*46*); screens with HIV-1 capsid mutants (*20*) and screens with whole genome libraries or targeted sublibraries with and without interferon (IFN) treatment (*17*, *26*). We invite contributions from groups using CRISPR-based functional genomics approaches to understand virus replication using other virus models to contribute data to CRISPRvirus.

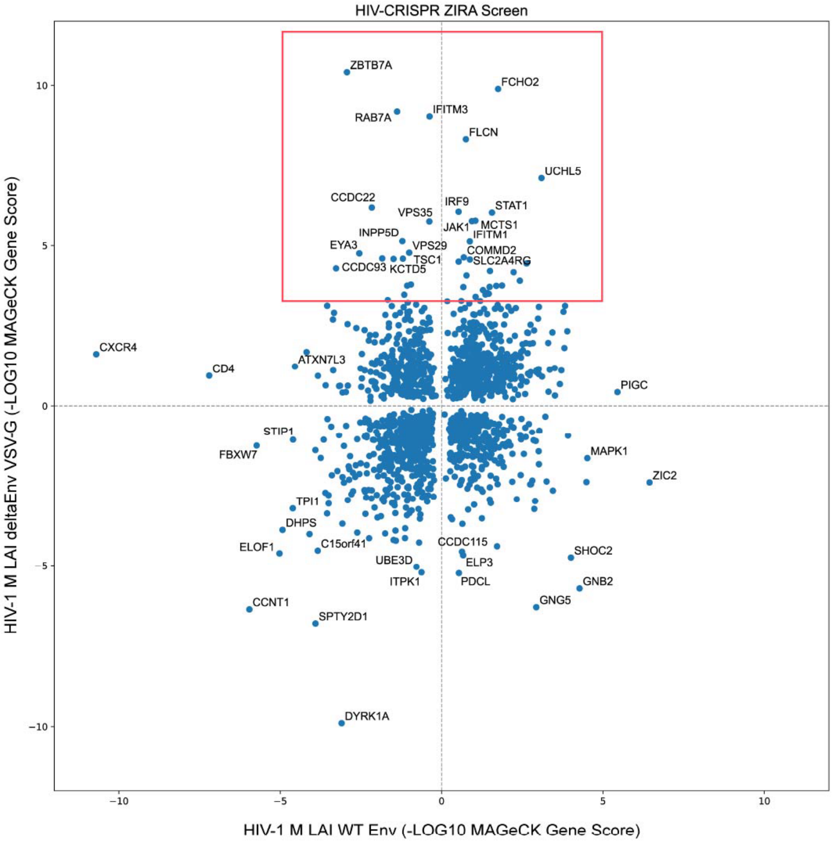


**Figure S4. Comparative ZIRA with native HIV Env and VSV-G.** Scatterplot showing -log10 MAGeCK Gene Scores from ZIRA sublibrary screening with HIV-1 M LAI (x-axis: native HIV env; y-axis: VSV-G pseudotyped) in THP-1 cells. Gene hits identified as specifically restricting VSV-G pseudotyped virus but not HIV-1 M LAI WT env virus are boxed in red.

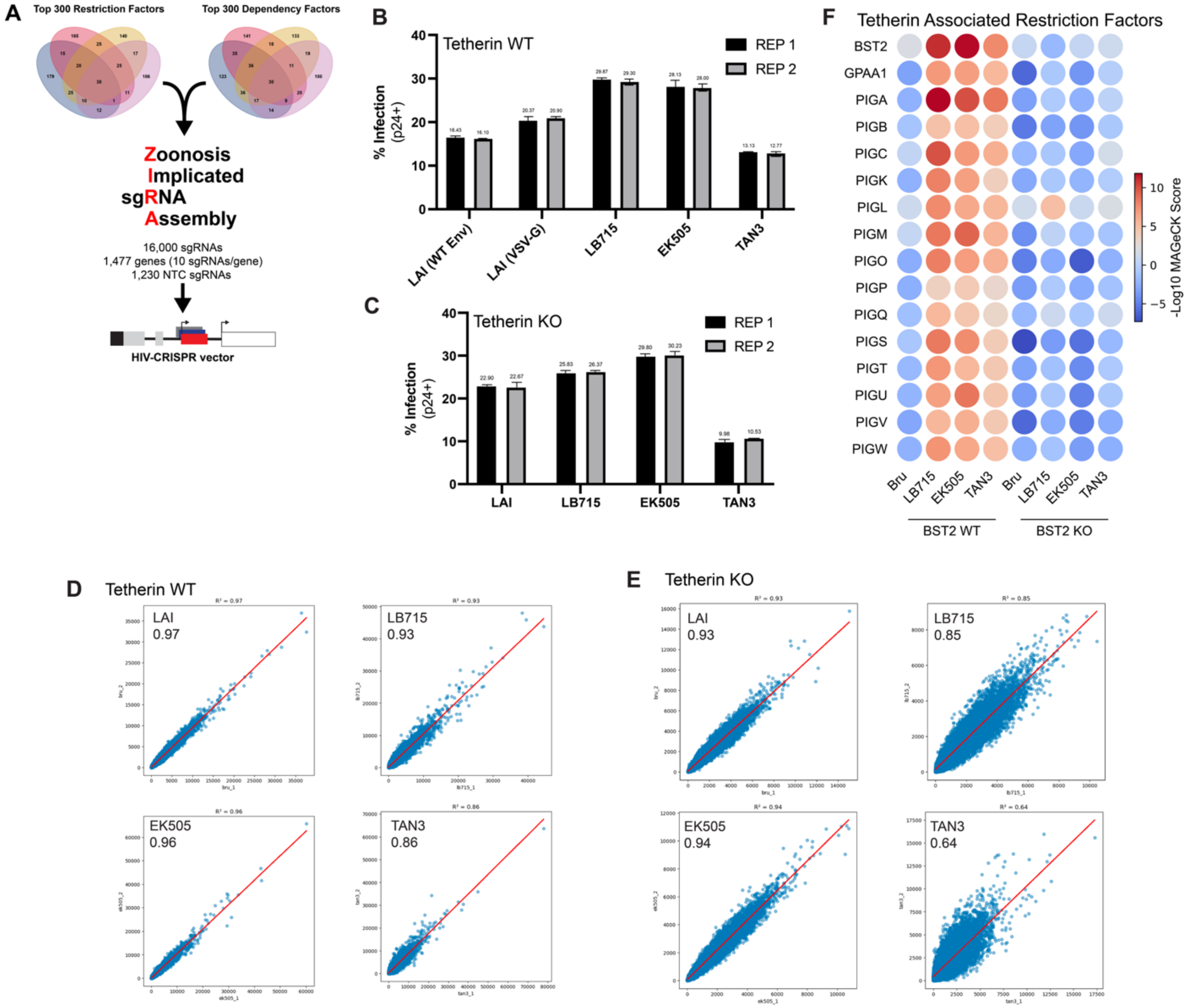


**Figure S5. ZIRA sublibrary screening.** (**A**) Overview of ZIRA sublibrary screening: selection of genes, synthesis and assembly of the Zoonosis Implicated sgRNA Assembly (ZIRA) into HIV-CRISPR. The top and bottom 300 genes from all whole-genome screens in Fig. 1 were selected, 10 guides were designed using CRISPick (*47*) to target each gene and the library as assembled into HIV-CRISPR to create the ZIRA HIV-CRISPR library. (**B** and **C**) Infection levels of ZIRA sublibrary HIV-CRISPR screens in wild-type (**B**) and Tetherin knockout THP-1 cells (**C**) across duplicate infections used in each screen. Infection measured by intracellular p24 antibody staining and flow cytometry. (**D** and **E**) Correlations of sgRNA counts between replicates for each ZIRA screen in wild-type (**D**) and Tetherin knockout (**E**) THP-1 cells. (**F**) Heat map comparing the -log10 MAGeCK gene scores for genes involved in Tetherin restriction and GPI-anchor biosynthesis between Tetherin (*BST2*)-expressing wild-type and Tetherin knockout THP-1 ZIRA screens for all four viruses.

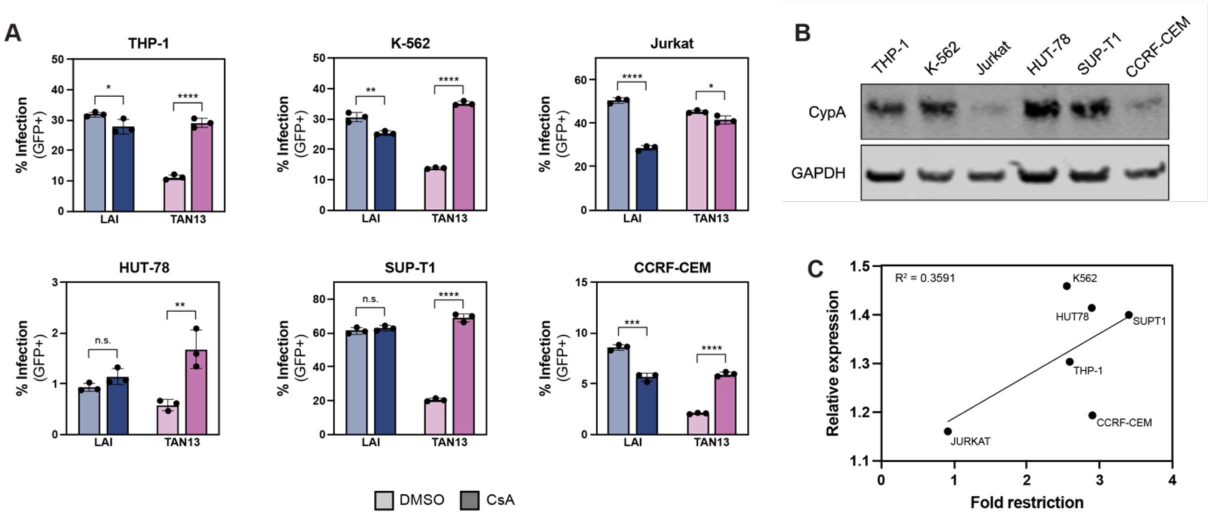


**Figure S6. CypA rescue infections across immune cell lines.** (**A**) Input-normalized infection of HIV-1 LAI and SIVcpz*Pts* TAN13 capsid chimera viruses in different immortalized lymphoid (Jurkat, HUT-78, SUP-T1, CCRF-CEM) and myeloid (THP-1, K-562) cell lines as measured by GFP fluorescence in cells treated with DMSO or CsA. (**B**) Immunoblot for CypA expression across cell lines tested in **A**. Blots were probed for GAPDH as a loading control. (**C**) Linear regression analysis showing the correlation between relative expression of CypA as quantified by immunoblot (y-axis) relative to fold restriction of SIVcpz*Pts* TAN13 capsid chimera virus by CypA (x-axis) across cell lines.

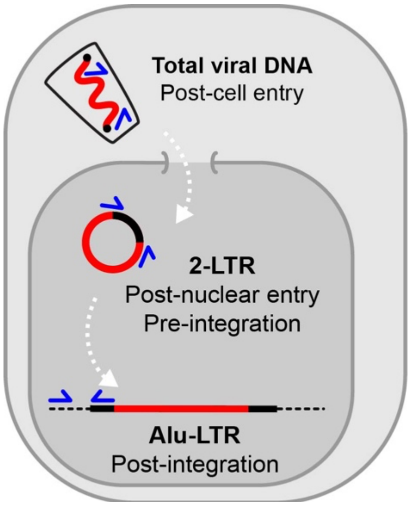


**Figure S7. HIV lifecycle qPCR assays.**
Schematic illustrating the viral cDNA products detected via qPCR-based lentivirus lifecycle assays. DNA is extracted from cells 24 hours after infection and amplified using three different primer sets (Total vDNA, 2'LTR and Alu-LTR). Total vDNA primers detect all viral DNA following reverse transcription. 2-LTR primers detect viral DNA products that form after nuclear entry. Alu-LTR primers detect integrated proviral DNA by amplification through binding to nearby repetitive genomic Alu elements. Primers are in Table S3.

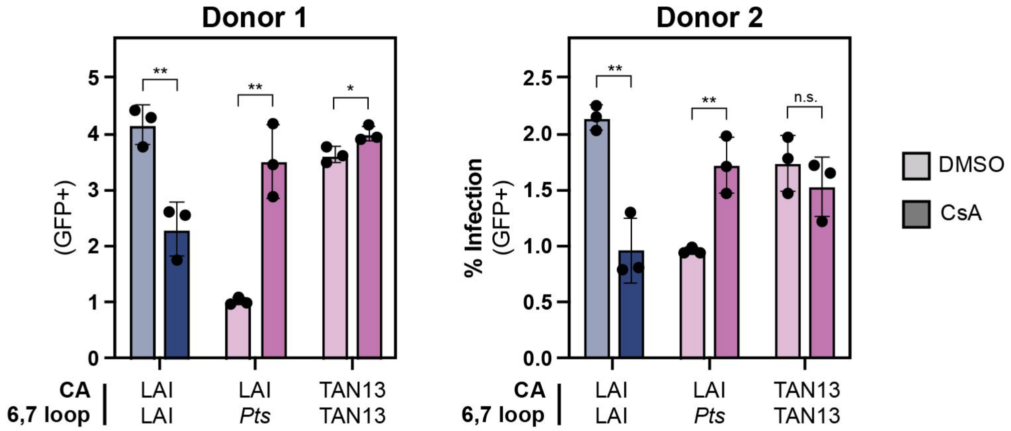


**Figure S8. CypA restriction of the SIVcpz*Pts* 6,7 loop in a primary CD4+ T cell infection model.**
Infection levels of HIV-1 LAI capsid (blue), LAI capsid chimera with the SIVcpz*Pts* 6,7 loop swap (purple) or SIVcpz*Pts* TAN13 capsid (purple) in DMSO or CsA (1 $\mu$M) treated primary CD4+ T-cells isolated from two healthy human donors. Data presented are triplicate infections from a single representative experiment. Experiments were performed three times with similar results. Statistical significance was determined by a two-tailed, unpaired Student's t-test between CsA treated or DMSO treated cells. *P<0.05; **P<0.01; ***P<0.001; ****P<0.0001; n.s.= not significant.

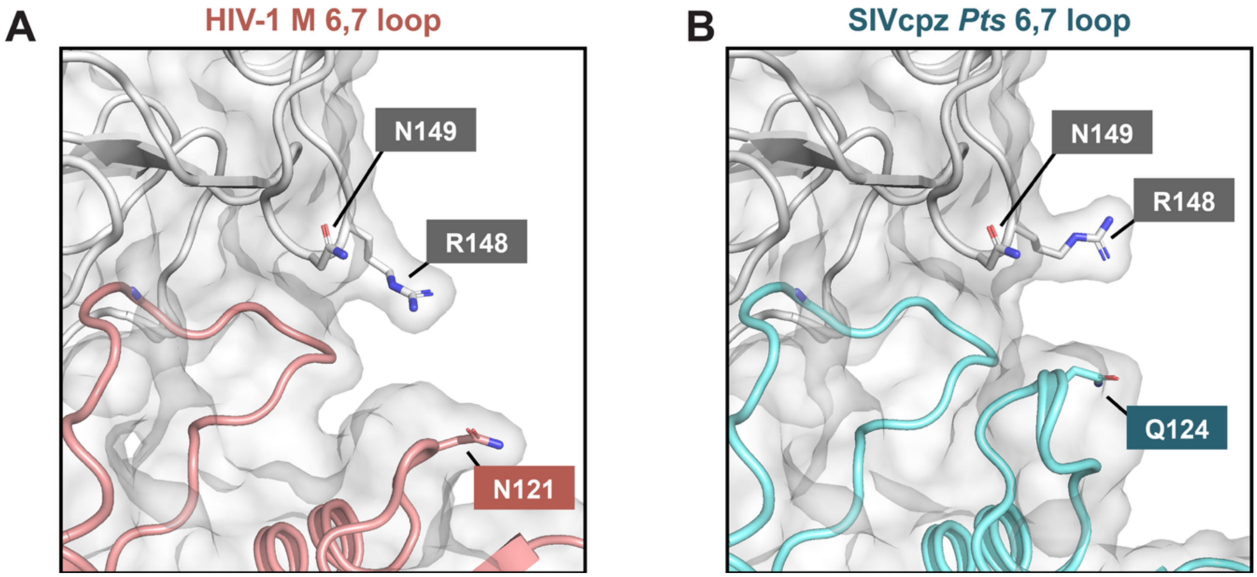


**Figure S9. CypA-capsid binding interfaces of HIV-1 M LAI capsid with WT or SIVcpz*Pts* 6,7 loop (no direct interaction with 6,7 loop).** X-ray crystal structures of the N-terminal domain of HIV-1 LAI CA (**A**) and HIV-1 LAI CA with the SIVcpz*Pts* 6,7 loop swap (**B**). Close-up views of the interface between capsid and CypA, highlighting residues at the interaction site shown in Fig. 4 (R148 and N149 in CypA; N121 and Q124 in capsid).

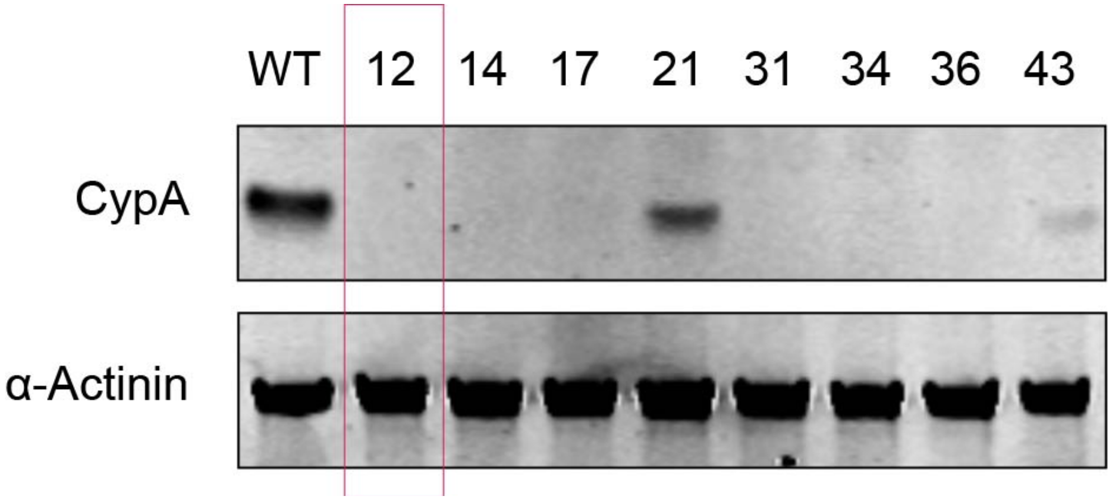


**Figure S10. Clonal CypA knockout THP-1 cell line.** Clonal knockout of CypA in THP-1 cells validated by immunoblot. Blots were probed for CypA and α-Actinin as a loading control. Clone boxed in red used in Fig. 4.

**A**

| Nucleotide | | Pos | Allele Frequency | | Impact |
|---|---|---|---|---|---|
| Ref | Alt | | *Ptt* | *Pts* | |
| T | C | 94 | 4/40 | 0/44 | H → R |
| T | C | 111 | 6/40 | 0/44 | K → R |
| C | T | 112 | 0/40 | 1/44 | V → I |
| G | A | 323 | 3/40 | 0/44 | P → L |
| C | T | 437 | 0/40 | 4/44 | R → H |
| G | A | 490 | 1/38 | 0/44 | S → L |
| G | A | -- | 1/40 | 0/44 | Splice variant |

Ref = Reference, Alt = Alternative, Pos = Amino Acid Position

**B**

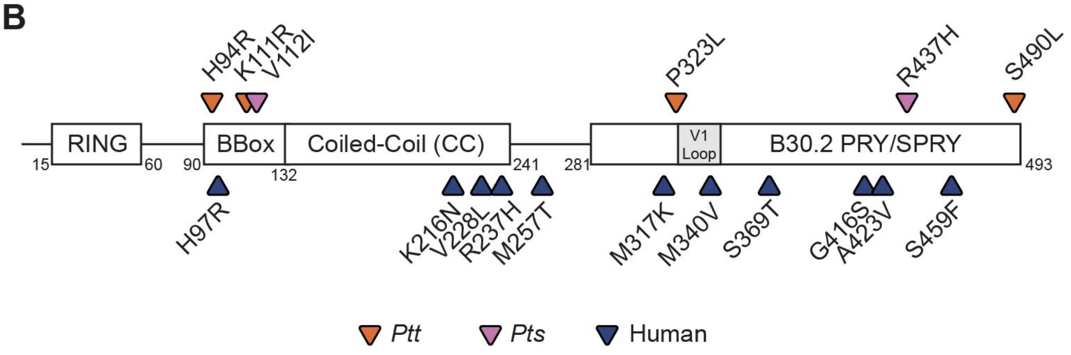


**Figure S11. TRIM5α genomic analysis.**
(**A**) Table of predicted high impact TRIM5α single nucleotide variants (SNVs) identified between Eastern (*Ptt)* and Central (*Pts*) subspecies of chimpanzee with indicated allele frequencies. (**B**) Visual representation of nonsynonymous (amino acid altering) TRIM5α SNVs mapped across functional domains (RING, BBox, Coiled-Coil) in Central (*Ptt* in orange) and Eastern (*Pts* in purple) chimpanzees. Nonsynonymous amino acid differences between reference chimpanzee and human TRIM5α orthologs are shown in blue.

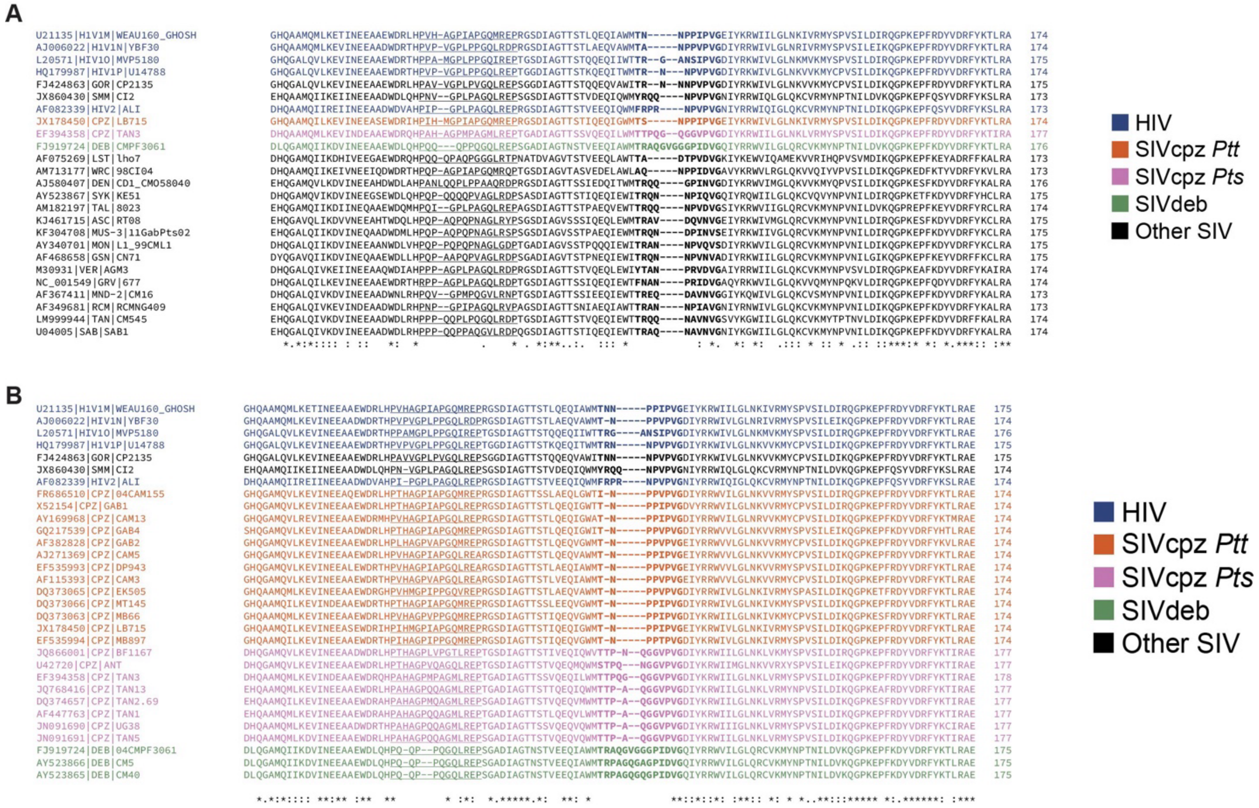


**Figure S12. HIV and SIV capsid alignment.** (**A and B**) Amino acid alignment using MAFTT of a portion of the capsid protein sequence across HIV and SIV sequences. HIV in blue; SIVcpz*Ptt* in orange; SIVcpz*Pts* in purple; SIVdeb in green; all other SIV in black. (**A**) Alignment of HIV-1 and a diverse sampling of a range of SIV capsids from a range of host species. (**B**) Alignment of HIV-1 along with a larger panel of zoonotic SIVcpz*Ptt*, nonzoonotic SIVcpz*Pts* and SIVdeb capsids. Dashes indicated alignment gaps. *, . and : indicate amino acid sequence conservation.
